# Spinal reflex modulation in the pelvic floor muscles through sensory stimulation from the lower limb

**DOI:** 10.64898/2026.06.09.730985

**Authors:** Yao Sun, Clare Cunningham, Jaynie F. Yang, E. Paul Zehr, Tania Lam

## Abstract

The pelvic floor muscles (PFM) are critical for maintaining continence and are a primary target of physiotherapy training to manage urinary incontinence. PFM training relies on voluntarily activating this muscle group, limiting its translation to neurological populations where recovery of bladder function is a priority. Indirect evidence suggests that sensory feedback from the lower limb can modulate PFM activity, which may provide alternative strategies for PFM training. Cutaneous reflexes have been used as a proxy to study how sensory inputs from the skin influence motoneuron excitability. To explore the feasibility of eliciting cutaneous reflexes in the PFM and their role in controlling PFM activity, this study examined: 1) the input-output relationship and 2) the nerve-specificity of PFM cutaneous reflex responses from tibial and superficial peroneal nerve stimulation. Twenty-one neurologically intact adults participated in this study. We recorded PFM and lower leg muscle electromyography while participants received cutaneous stimulation to the right distal tibial nerve, bilateral distal tibial nerve, or right superficial peroneal nerve in a standing position. We delivered stimulation at the intensity below motor threshold (MT), 1.2 x MT and 1.5 x MT and quantified tibial-PFM reflex amplitude over a 50-150 ms window after stimulation. PFM reflex responses were evoked from both nerves’ stimulation. Reflex amplitude increased with stimulus intensity with tibial nerve stimulation but not with superficial peroneal nerve stimulation. Bilateral tibial nerve stimulation evoked larger responses compared to unilateral stimulation. These findings support the existence of neural connections between lower limb afferents and the PFM, and open up possibilities for designing rehabilitation strategies to manage pelvic health conditions in people with neurological disorders.

**New & Noteworthy:**

- Cutaneous sensory feedback from the foot, specifically that related to limb loading, can evoke reflex responses in the pelvic floor muscles
- Nerve–specific modulation was observed. Reflex amplitudes in the pelvic floor muscles increased with tibial nerve stimulation intensity but not with superficial peroneal nerve stimulation.

## Introduction

The pelvic floor muscles (PFM) are a network of muscles and connective tissue at the base of the abdominal cavity (Bø 2004). Importantly, these muscles are anatomically continuous and activated synergistically with the urethral and anal sphincters (Jorge and Bustamante-Lopez 2022). Thus, activation of the PFM functions to provide mechanical support of the pelvic organs as well as to ensure continence during elevated intra-abdominal pressure (IAP), such as during coughing or sneezing (Meagher et al. 1993). Other full-body activities, such as jogging or jumping, also increase IAP and engage the PFM (Grillner et al. 1978; Moser et al. 2018; Stafford et al. 2012). In these scenarios, the elevated IAP is accompanied by somatosensory inputs from the lower limb due to changes in posture and lower body loading, highlighting a potential interaction between the neural control of the PFM and lower limb muscles.

There is anatomical and physiological evidence that somatosensory inputs from the leg can have a modulatory effect on the muscular structures that control urinary function. The spinal cord circuits mediating lower urinary tract function, as well as bowel and sexual functions, overlap with those controlling sensorimotor function from the lower leg. The pudendal nerve, which innervates the PFM, originates from the S2-S4 spinal segments, and the tibial and peroneal nerves, which innervate the lower leg, originate from L4 to S3. This anatomical overlap at the level of the lumbosacral spinal cord supports the possibility of spinal neuronal circuits mediating the interaction between lower limb sensory inputs and structures supporting urinary function. Further, data from animal studies show that inputs from hindlimb afferents can impact bladder physiology. In uninjured, anesthetized cats, electrical stimulation of the tibial nerve, which innervates the plantar surface of the hindpaw, is effective in reducing the pressure and increasing the capacity of a filled bladder (Tai et al. 2011a). These effects were not observed when stimulation was applied to the forelimb, confirming that the bladder and its associated neuromuscular structure specifically responded to the sensory inputs from the hindlimb (Tai et al. 2011a). It is also notable that a long-standing clinical approach for overactive bladder, with roots in Chinese traditional medicine, is electrical stimulation of the distal tibial nerve at the ankle (Booth et al. 2018; Wall and Heesakkers 2017). Several randomized controlled trials confirm that 6 to 12 sessions of distal tibial nerve stimulation can be effective for improving symptoms of overactive bladder (Finazzi-Agrò et al. 2010; Lashin et al. 2021; Peters et al. 2010). Despite the promising functional changes observed in animal models and clinical trials, the specific mechanisms of how somatosensory inputs from the lower limb affect the PFM are not clear in human participants.

To understand the neural circuits mediating the effects of somatosensory input on motor neuron output, cutaneous reflexes, obtained by measuring the muscle response following afferent nerve stimulation, are commonly used (Zehr 2006; Zehr and Stein 1999). The characteristics of reflex modulation observed in the tested muscles can reveal how the nervous system utilizes sensory inputs from the skin to modulate muscle activity and control movement. For example, both animal and human studies have shown that sensory inputs from different nerves around the feet differentially modulate the excitability of various leg muscles during walking (Pearcey and Zehr 2019; Zehr et al. 1998a). This nerve-specific modulation was also reported for bladder function in the cat model; in contrast to the increased bladder capacity following tibial nerve stimulation mentioned above, electrical stimulation of the superficial peroneal nerve (which innervates the top of the foot) has the opposing effect of decreasing bladder capacity and facilitating voiding (Chen et al. 2022; Zhao et al. 2020). These results suggest that sensory inputs from different nerves of the hindlimb may also affect the muscular structures controlling lower urinary tract function differentially.

There is scant research investigating how sensory inputs around the lower limb modulate PFM activity in human participants. Pedersen et al. (Pedersen et al. 1978) reported that tibial nerve stimulation evoked a reflex response in the external anal sphincters in adults with and without neurological lesions. Hallet et al. (Hallett et al. 1984) also reported reflex responses in the urethral sphincters in children with spinal cord lesions following pinprick of the foot sole. These two studies not only suggested a neural connection between the lower limb afferent and pelvic structure at the lumbosacral level but also the potential of using cutaneous reflexes as a proxy to study control mechanisms of PFM activity in human participants.

To explore the feasibility of evoking cutaneous reflexes in the PFM and using it as a probe for understanding the modulatory role of somatosensory inputs from lower limb on PFM activity, this study investigated: 1) PFM responses following stimulation of the distal tibial and superficial peroneal nerve at different intensities; and 2) whether PFM respond differently to distal tibial and superficial peroneal nerve stimulation.

## Methods

### Participants

We enrolled 21 healthy, uninjured adults (10 females, mean ± standard deviation age: 25.4 ± 4.6 years old, height: 171.7 ± 8.3 cm, weight: 74.5±16.3 kg) to this study. All participants were free of any clinical condition affecting bladder, bowel, or sexual function. Participants were excluded if they were pregnant, or had been pregnant within the past 6 months, had ever undergone surgery involving the sex organs or perineum (excluding circumcision in males as long as the surgery was at least 6 months previous), had a genital piercing, or had been diagnosed with any pelvic pain disorder (e.g. vaginismus, vulvodynia). The study protocol was approved by the University of British Columbia UBC Clinical Research Ethics Board (Protocol number: H24-00251). All participants provided their voluntary, informed consent prior to data collection.

### Electromyography (EMG)

Following skin preparation with alcohol swabs and NuPrep (DO-NP Series, Aurora, USA), we affixed surface electrodes (Trigno, Delsys Inc., Boston, USA) on the skin overlying the soleus (SOL), tibialis anterior (TA), gluteus maximus (GM) and rectus abdominis (RA) on the right side. To record PFM EMG, a pair of disposable electrodes (Impulse Medical Technologies, Seattle, WA) was affixed approximately 1cm bilaterally from the anus and then connected to snap lead Delsys Trigno Sensors (Delsys Incorporated, Natick, USA). For the PFM EMG set-up, we provided participants with detailed instructions and a diagram to allow them to affix PFM electrodes on themselves in a private setting. We confirmed that PFM electrodes were placed correctly before data collection by asking participants to voluntarily contract their PFM or cough. If a clear PFM activity was seen in the data collection systems without coactivation from GM, we confirmed the electrodes were placed correctly. All EMG data were collected at a sampling rate of 2000 Hz using a customized LabVIEW program (National Instruments, Austin, USA).

### Electrical stimulation

We placed surface electrodes (Thought Technology Ltd., Quebec, Canada) on participants’ distal tibial nerve bilaterally and the superficial peroneal nerve of the right foot (Figure 1). Cutaneous stimulation was delivered using a custom LabVIEW program and a GRASS 88 stimulator with SIU stimulus isolation and CCU1 constant current unit (Grass Instruments, Astro-Med, West Warwick, USA).

**Figure 1.**
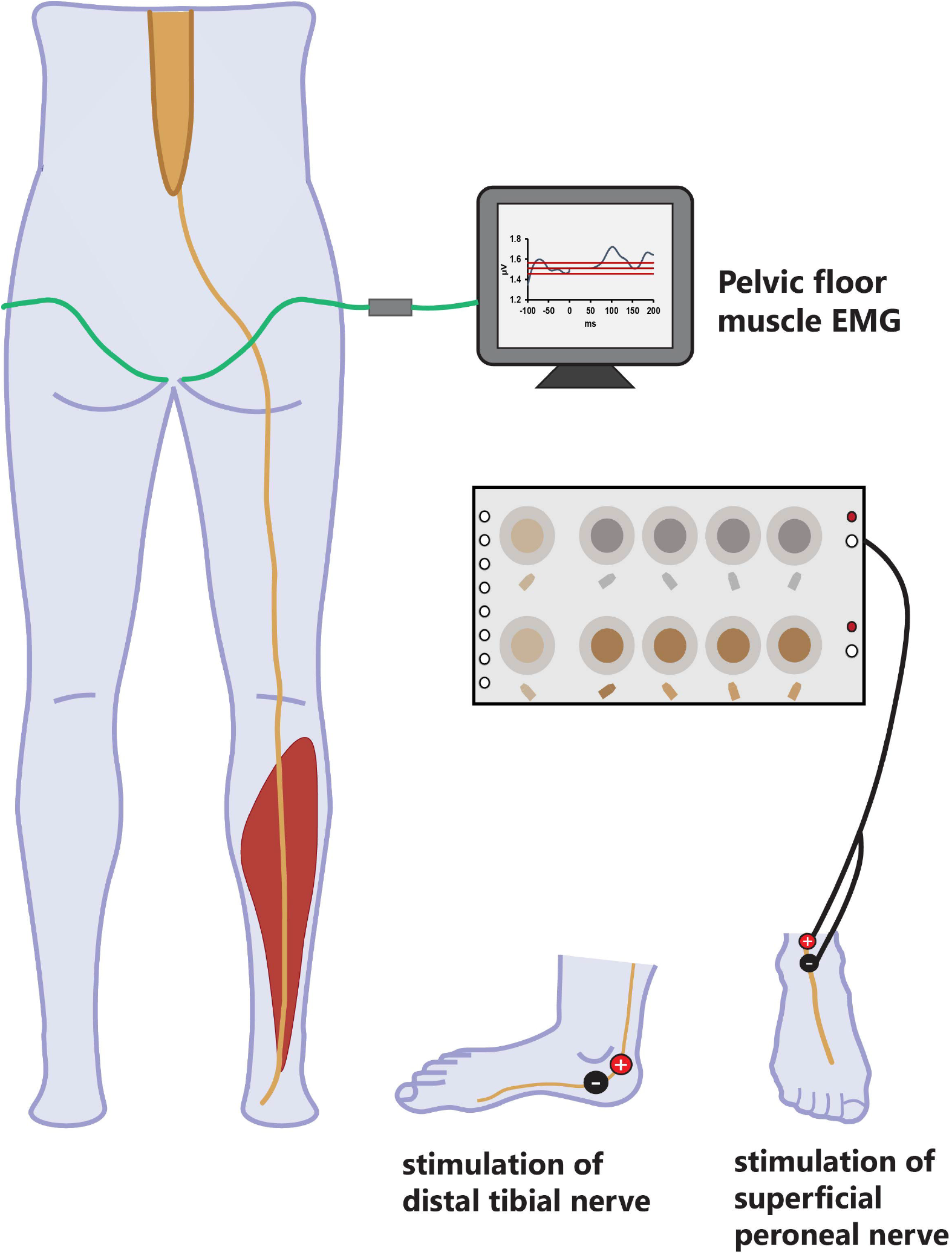
Experimental set-up

Prior to data collection, we first determined the perceptual threshold (PT), radiating threshold (RT), and motor threshold (MT) of each nerve. PT was defined as the minimum current required to evoke a detectable tactile sensation. RT was defined as the minimum current that generates a clear radiating sensation of the nerve’s innervation area. MT was defined as the minimum current that evoked a toe twitch. Each threshold measurement was repeated two to three times for consistency.

Stimulation was delivered as a train of five 1-ms pulses at 300 Hz, following previous studies examining cutaneous reflex responses in the leg muscles (Pearcey and Zehr 2020; Zehr and Chua 2000; Zehr et al. 2001). The parameters were chosen to create a strong sensation along the plantar (via distal tibial nerve stimulation) or dorsal surface (via superficial peroneal nerve stimulation) of the foot, which has been described as “buzzing”, “vibrating” and/or “tingling”. To understand the intensity-response relationship of the PFM, we recorded responses at three stimulus intensities: just below the MT (<MT), 1.2 x MT, and 1.5 x MT for each nerve.

### Experimental procedure

A schematic diagram of the experimental set-up and procedure is presented in Figure 1. Data collection was completed with participants standing with feet shoulder-width apart. To determine PFM activity during attempted maximum voluntary contraction (aMVC), we provided verbal instructions to guide participants to contract their PFM without co-activating other muscles. Real-time EMG signals were displayed on a computer monitor as visual feedback. Once the participants were confident in producing PFM contractions without co-contraction of the other muscles, we recorded the EMG during aMVC from two attempts. Then, the 3 nerve-stimulation conditions were completed with stimuli delivered: 1) unilaterally to the right tibial nerve only (Tib-R), 2) bilaterally to the left and right tibial nerves (Tib-B), and 3) unilaterally to the right superficial peroneal nerve only (SP-R). For each nerve-stimulation condition, we delivered 100 stimuli for each stimulation intensity (<MT, 1.2 x MT, and 1.5 x MT) with random interstimulus intervals between 2-3 s. Participants were allowed to sit and rest between trials.

Before the start of each condition, we delivered one to two stimuli to the participants to make sure the intensity did not exceed their pain thresholds. None of the participants reported pain during the <MT trials for any condition. Stimulus intensity had to be adjusted in some participants in the SP-R condition only. Five participants reported pain when the SP-R stimulus was delivered at 1.2 x MT. In these participants, we replaced the 1.2 x MT and 1.5 x MT conditions with 1.5 x RT, and just below the pain threshold to maintain three distinct levels of stimulation below the pain threshold. One additional participant reported pain with SP-R at 1.5 x MT stimulation, so we reduced the stimulation intensity to just below pain threshold (1.4 x MT) for this participant.

### Data analysis

Data were analyzed offline using custom-written MATLAB programs (Version R2023a, The Mathworks, Natick, USA). After offset and gain correction, the EMG signal was rectified and low-pass filtered at 30 Hz with a 4th-order dual-pass Butterworth filter.

To calculate the EMG amplitude during the aMVC trials, the relaxation phase before contraction and during the maximal contraction phase were selected manually. The average of a 0.5-s window around the minimum of the relaxation phase and the maximum during contraction was calculated; the difference between the averaged maximum and minimum was defined as the aMVC EMG. The mean of the two aMVCs was used to normalize the EMG amplitudes for each participant.

Before characterizing the reflex response in PFM, we first measured the pre-stimulus (background) EMG to ensure each stimulus was applied during stable background muscle activity. The mean background EMG from 75ms to 5ms before stimulus onset was calculated for each trial. Then, the mean and standard deviation of the background EMG across the 100 trials within each condition were calculated for each participant. Trials with background EMG falling outside 2 standard deviations of the grand mean were removed from further analysis.

PFM reflex responses were characterized by three different measures: 1) the number of conditions that evoked a PFM reflex; 2) the onset and peak latencies of the PFM reflex response, and 3) the amplitude of net reflex responses. To determine if there is a PFM reflex response evoked in each condition, trials with stable background EMG were averaged. The post-stimulus EMG was compared with the averaged background EMG. If the post-stimulus EMG amplitude exceeded the mean ± 2 standard deviation of the background at any time within a 150ms window post-stimulation, we defined that a reflex response was present in that condition. The time when the post-stimulus EMG exceeded the mean ± 2 standard deviation of the background EMG was defined as the reflex onset latency, and the time when the post-stimulus EMG reached its highest value was defined as the peak reflex latency.

The PFM reflex amplitude for each trial was quantified as the relative change from the background activity. We first calculated the mean EMG amplitude over the 50-150ms window post-stimulus as the net response (Zehr et al. 1997). The difference between this post-stimulus net response and the background EMG was expressed as a % of the background EMG. The amplitude of the net reflex response was calculated for all the trials that with stable background EMG, regardless of whether a PFM reflex response was evoked in that conditions. This would allow us to use the Linear Mixed Effect Model to examine the potential effects of stimulation intensity and background EMG (more details in the next section)

To confirm that the responses observed in the PFM were not from crosstalk of a synergistic or nearby muscles (i.e. GM and RA), we examined the EMG of GM and RA in each condition. No visible responses were observed in the RA muscle in any participant, but increased post-stimulus GM muscle activity was observed in some participants (e.g. Fig. 2B). The presence of GM responses were determined in the same ways as PFM. For conditions that presented with responses in both the PFM and GM, the latency of the peak reflex response was calculated for further analysis.

**Figure 2.**
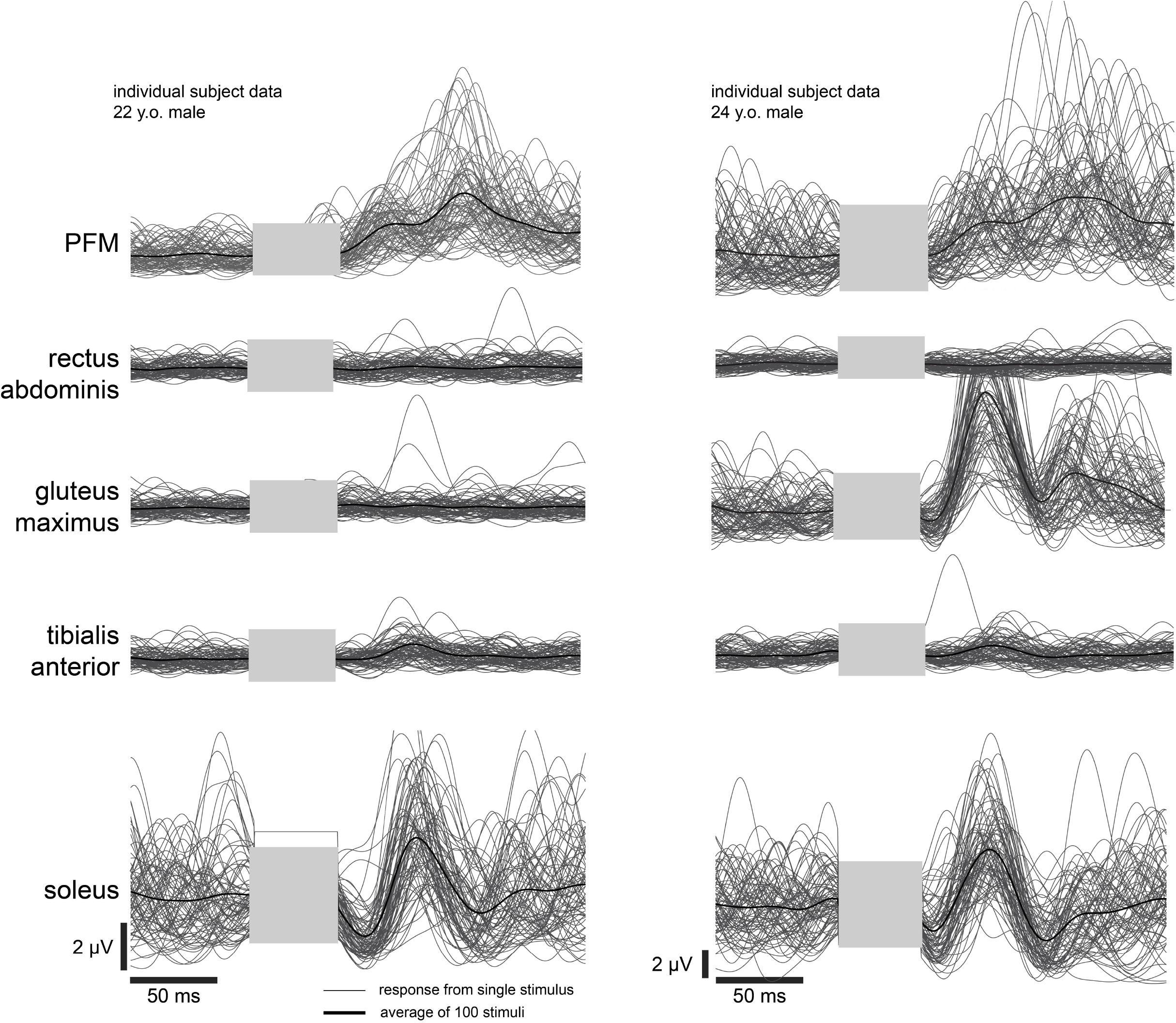
Reflex response of pelvic floor muscle (PFM) and other muscles from two individual participants following bilateral tibial stimulation at the intensity of 1.5 x MT. The thin grey line represents reflex response from one stimulus. Thick black lines represent the averaged reflex response from 100 stimuli. The grey box represents the removed stimulation artifact.

## Statistical analysis

To confirm stability of the background EMG across trials within each participant, we performed two-way repeated-measures ANOVA with main effects of Nerve (Tib-R, Tib-B, SP-R) and Intensity (<MT, 1.2 x MT, 1.5 x MT) on the mean background EMG of each trial using SPSS (IBM SPSS Statistics 30.0; Chicago, IL).

To characterize the modulation of PFM reflex responses with increased sensory inputs from the bottom of the feet (Tib-R vs Tib-B), and with sensory inputs from different nerves (Tib-R vs SP-P), we used the lme4 package in RStudio (version 2024.04.2) to create linear mixed models (LMM) for data analysis. Reflex amplitudes calculated from individual trials in each condition were fitted with LMM models with fixed effects of Nerve, Intensity, background EMG, and random effects of Participant. Reference condition was set as Tib-R for the effects of Nerve, and <MT for the effects of Intensity. As the effect of Intensity may vary by the nerve(s) being stimulated or across participants, we also explored the interaction effects of Nerve x Intensity and the random effects with or without a random slope of Intensity. We identified the best fitting model based on a combination of Akaike Information Criterion (AIC), the Likelihood Ratio Test (LRT) and the Marginal R^2^.

To examine if responses observed in the PFM were independent of responses in GM, we used a grid plot to examine for the presence of PFM responses corresponding with the presence of a response in the GM in each condition across participants. When corresponding responses occurred in the two muscles, peak response latencies from these muscles were compared using Spearman’s correlation coefficient. Statistical significance of all tests was set at P ≤ 0.05.

## Results

All participants completed all trials in Tib-R and Tib-B conditions. The SP-R condition was missing from one participant due to time constraints during data collection. Across all conditions, 72 to 95 out of 100 trials with stable background EMG were used for further analysis. Repeated measure ANOVA on these data confirmed that background EMG did not differ by Nerve (F(2,38) =2.669, p=0.082) or Intensity (F(2,38) =0.786, p=0.463).

### Characteristics of PFM reflex responses

PFM reflex responses were present in 160 out of a total of 188 nerve-intensity conditions (21 participants x 3 nerve-stimulation conditions x 3 intensities). With an averaged reflex onset latency at 54.5 ± 16.8ms and peak latency at 113.0 ± 19.1ms. Different from cutaneous reflexes observed in the leg or arm muscles, where cutaneous reflexes often show distinct early and middle latency responses, an overall facilitatory response was observed in the PFM in a 50 - 150 ms window after stimulation.

Reflex responses were observed in the GM muscle in a few participants after either tibial or SP nerve stimulation. Figure 3 summarizes the presence of PFM and GM responses in each condition for each participant. In 70 out of 188 nerve-intensity conditions, responses were present in both PFM and GM muscles (black cells in Figure 5). Spearman correlation indicated the latencies of the peak responses in the PFM and GM in these cases were not correlated (p=0.32, Fig. 4), suggesting most of the responses observed in PFM were likely not from cross-talk from GM. However, 10 trials show the GM reflex reaches its peak just before PFM peak reflex with the difference of peak reflex latency less than 4ms (i.e. smaller than the typical action potential delay at one synapse). We could not completely rule out the influence of GM response on the PFM reflex in these 10 trials. But considering these trials were from different conditions among different participants. The GM response may reflect random individual variance instead of a confounding effect.

**Figure 3.**
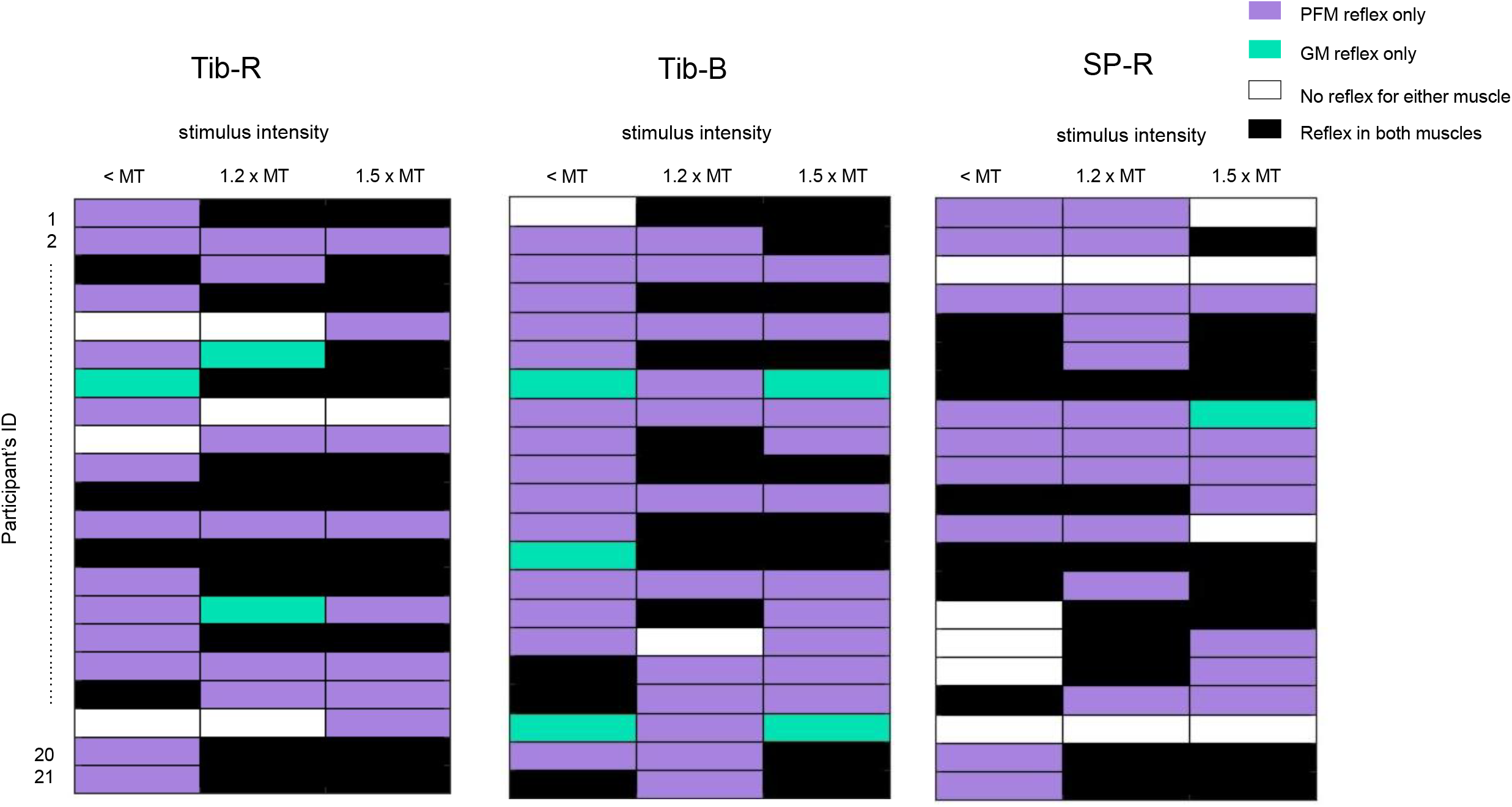
Summary of reflex appearance in pelvic floor muscles (PFM) and gluteus maximus (GM). In each checkboard figure, rows represent the results from 21 participants, and columns represent 3 stimulation intensities when stimulation was applied to the right tibial nerve (Tib-R), bilateral tibial nerve (Tib-B) and right superficial peroneal nerve (SP-R). Purple, green, black and white squares represent a reflex was observed in pelvic floor muscles (PFM) only, gluteus maximus (GM) only, both muscles, or not observed in either muscle, respectively.

**Figure 4.**
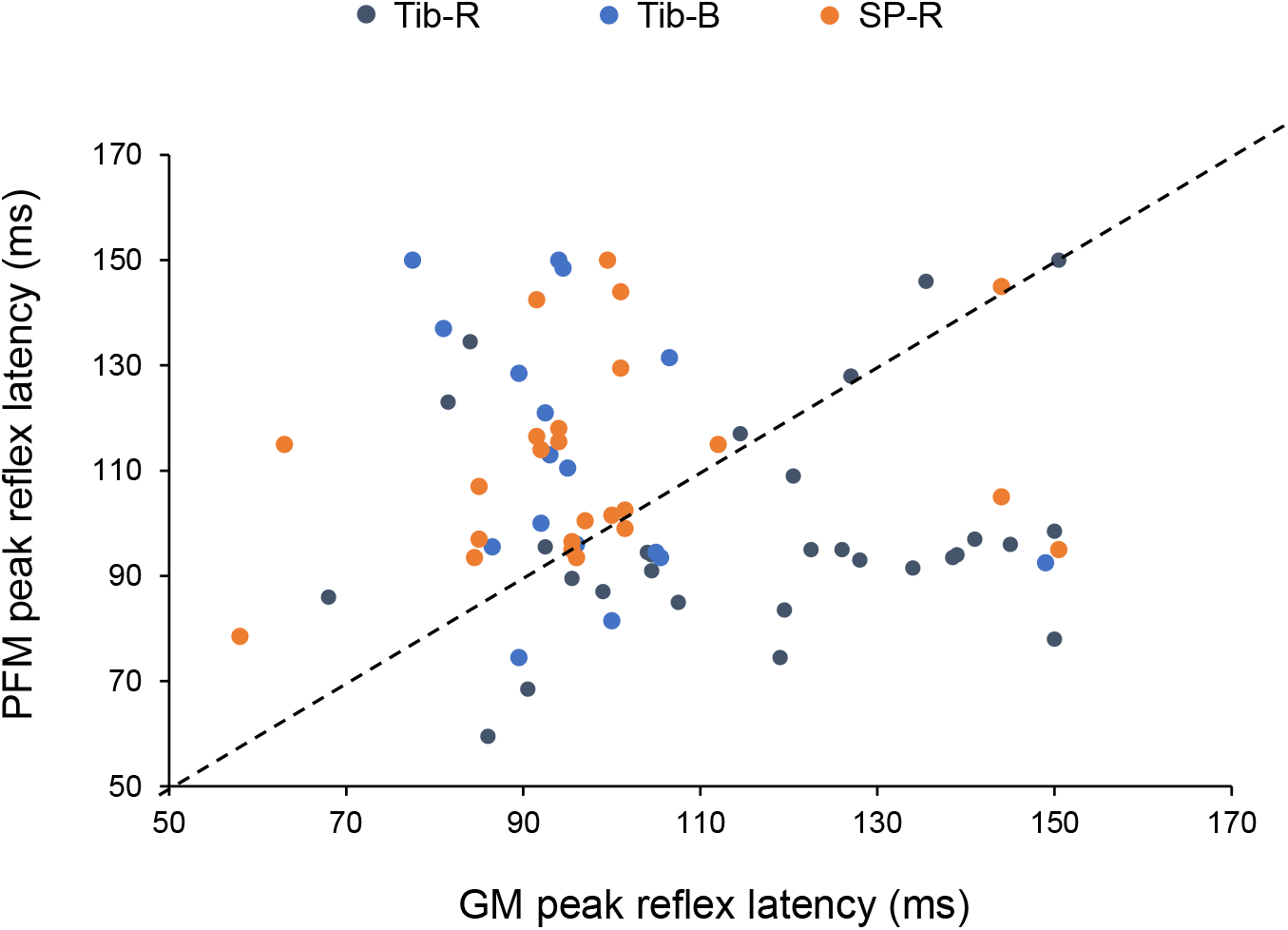
Comparison of peak reflex latency between pelvic floor muscle (PFM, y-axis) and gluteus maximus (GM, x-axis). The diagonal dashed line represents

**Figure 5.**
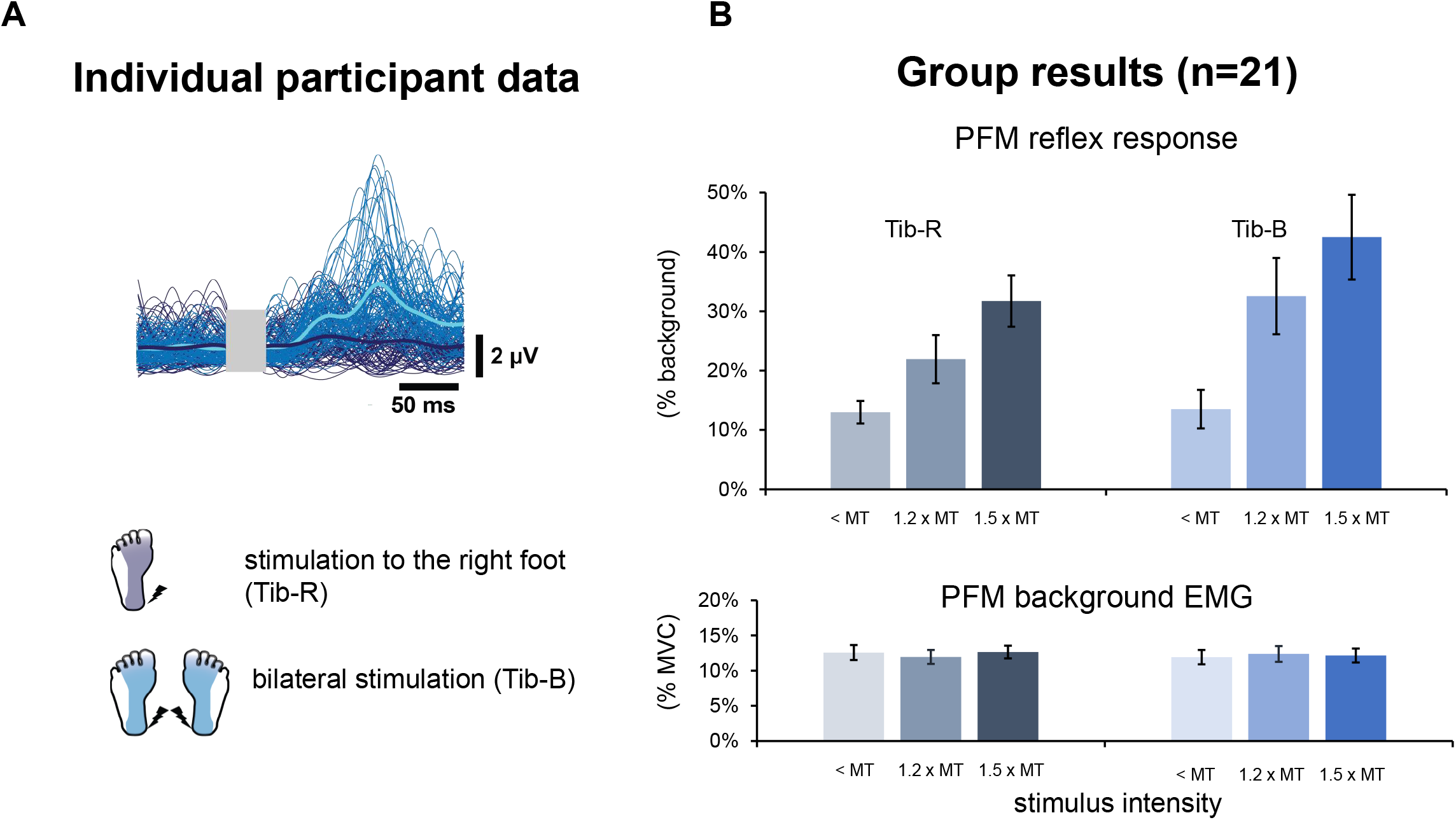
Reflex response in pelvic floor muscle (PFM) following tibial nerve stimulation to the right foot (Tib-R) and to both feet (Tib-B). **A.** One individual participant’s data. Data were collected from tibial nerve stimulation on the right side (dark blue) and bilateral stimulation (light blue) with the simulation intensity at 1.5 x motor threshold (MT). Thin lines represent the response from one stimulus; thick lines represent the average from the same trial. The dermatome of the tibial nerve is shown as shaded areas in the diagram at the bottom. **B**. Group results. The x-axis represents stimulation intensity at below, 1.2 x, and 1.5 x MT. Error bars represent the standard error of the mean across participants.

**Figure 6.**
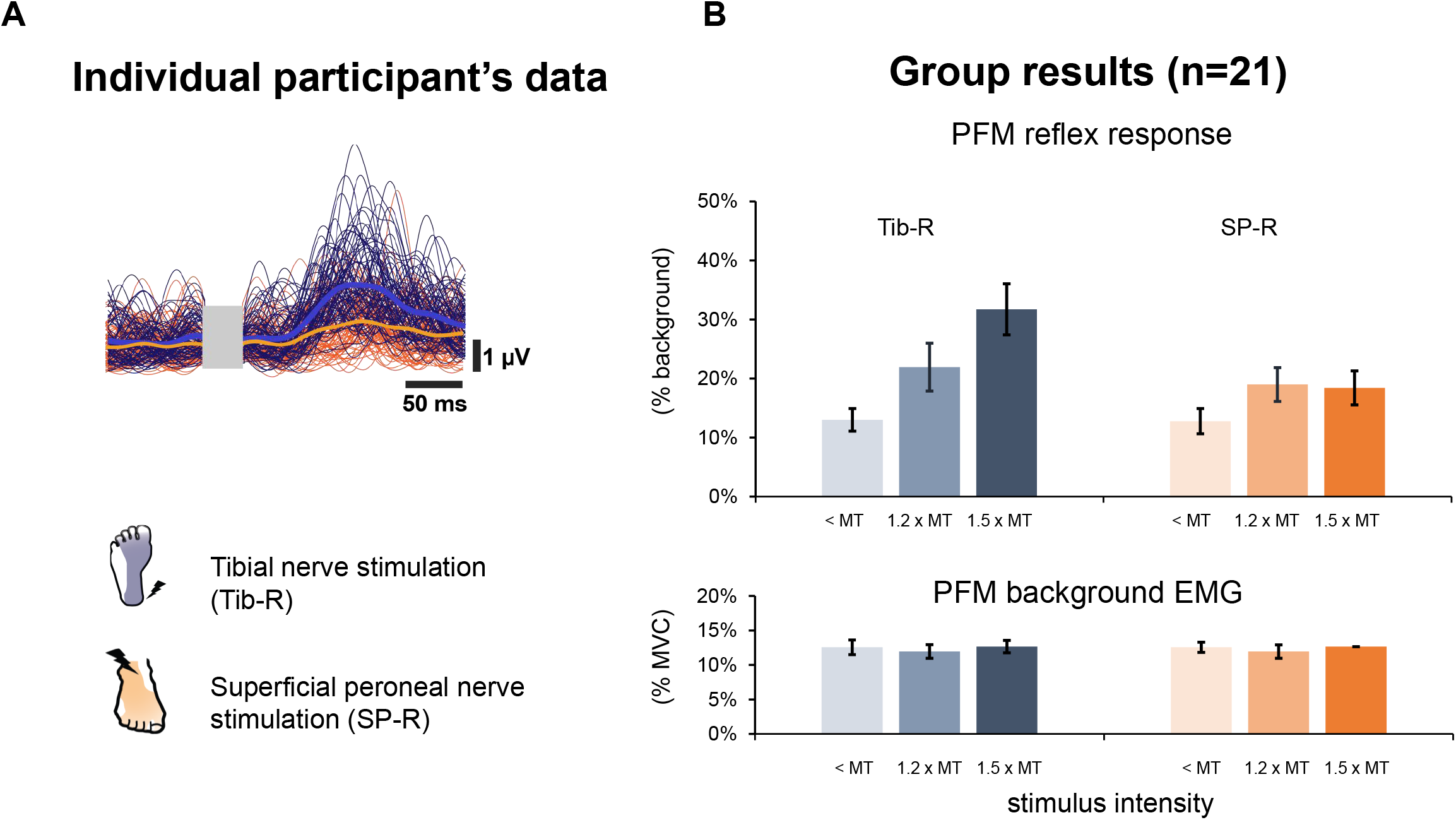
Reflex response in pelvic floor muscle (PFM) following tibial nerve stimulation (Tib-R) and superficial peroneal stimulation (SP-R) to the right foot. **A.** One individual participant’s data. Data were collected from tibial nerve stimulation (dark blue) and superficial peroneal nerve stimulation (orange) with the simulation intensity at 1.5 x motor threshold (MT). Thin lines represent the response from one stimulus; thick lines represent the average from the same trial. The dermatome of the tibial nerve and superficial peroneal nerve is shown as shaded areas in the diagram at the bottom. **B**. Group results. The x-axis represents stimulation intensity at below, 1.2 x, and 1.5 x MT. Error bars represent the standard error of the mean across participants.

### Nerve-specific and stimulus intensity effects

The LMM that included the interaction effect of Nerve x Intensity and random slope of Intensity significantly improved model fit based on the log likelihood ratio test (χ^2^ (4) = 263.9, p < 0.001) and Marginal R^2^ = 0.231 and Conditional R^2^ = 0.396. This model revealed significant effects of Intensity (p <0.001), Nerve (p<0.001), background EMG (p <0.001), as well as significant interaction effects of Nerve x Intensity (p<0.001) on PFM reflex amplitudes. Specifically, compared to the Tib-R condition, Tib-B had significantly larger PFM reflex responses at the intensity of 1.2 x MT (estimated difference = 0.15, 95%CI [0.11, 0.19], p < 0.001) and 1.5 x MT (estimated difference = 0.11, 95%CI [0.07, 0.15], p < 0.001, Figure 4). SP-R evoked smaller PFM reflex responses at <MT (estimated difference = -0.08, 95%CI [-0.11, -0.05], p < 0.001) and 1.5 x MT stimulation (estimated difference = -0.19, 95%CI [-0.23, -0.15], p < 0.001) compared to Tib-R (Figure 5, Table 1).

**Table 1.**
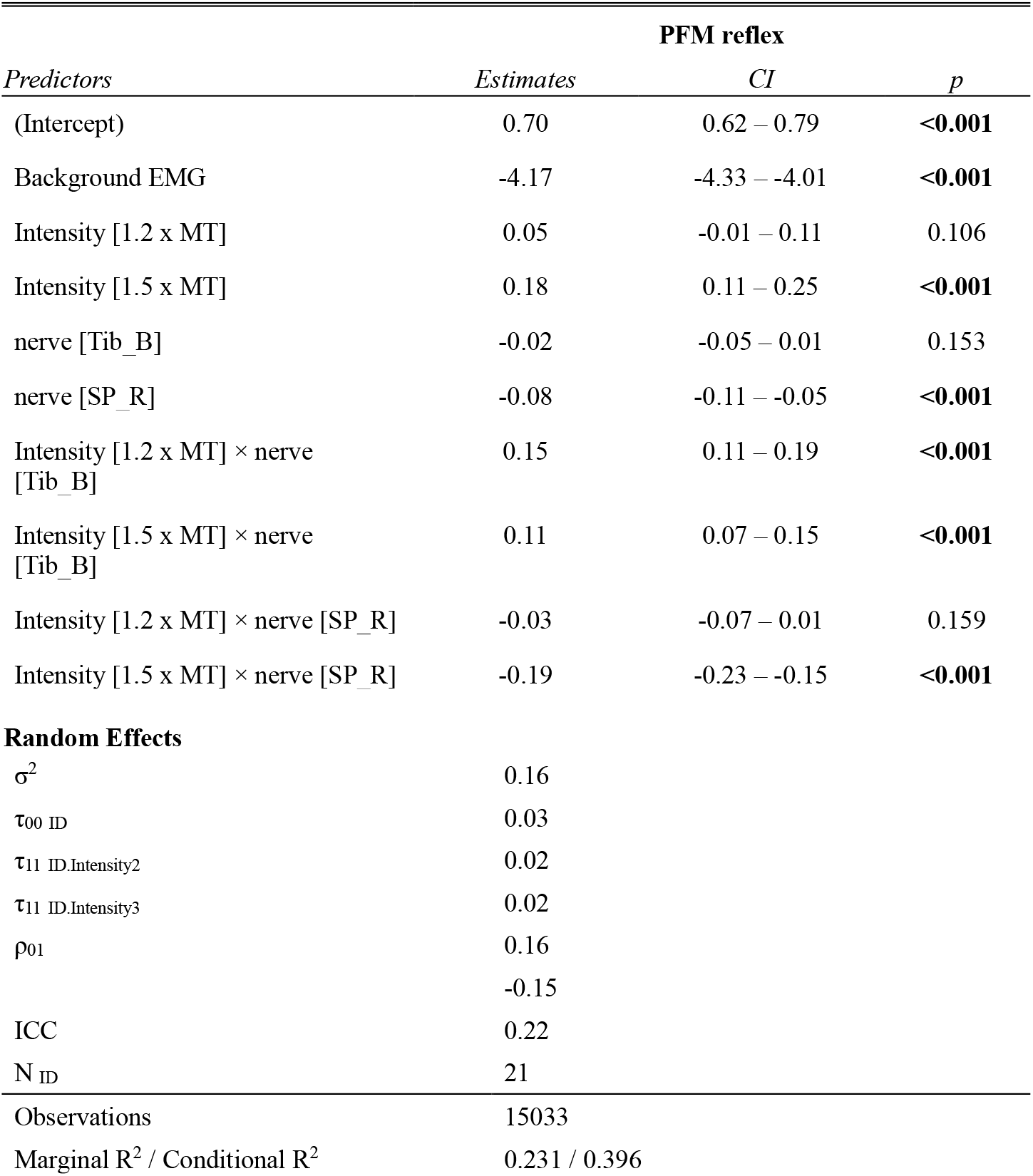
Linear mixed model analysis results of the PFM reflex.

| *Predictors* | **PFM reflex** |  |  |
| --- | --- | --- | --- |
|  | *Estimates* | *CI* | *p* |
| (Intercept) | 0.70 | 0.62 – 0.79 | **<0.001** |
| Background EMG | -4.17 | -4.33 – -4.01 | **<0.001** |
| Intensity [1.2 x MT] | 0.05 | -0.01 – 0.11 | 0.106 |
| Intensity [1.5 x MT] | 0.18 | 0.11 – 0.25 | **<0.001** |
| nerve [Tib_B] | -0.02 | -0.05 – 0.01 | 0.153 |
| nerve [SP_R] | -0.08 | -0.11 – -0.05 | **<0.001** |
| Intensity [1.2 x MT] × nerve [Tib_B] | 0.15 | 0.11 – 0.19 | **<0.001** |
| Intensity [1.5 x MT] × nerve [Tib_B] | 0.11 | 0.07 – 0.15 | **<0.001** |
| Intensity [1.2 x MT] × nerve [SP_R] | -0.03 | -0.07 – 0.01 | 0.159 |
| Intensity [1.5 x MT] × nerve [SP_R] | -0.19 | -0.23 – -0.15 | **<0.001** |
| **Random Effects** |  |  |  |
| $\sigma^2$ | 0.16 | | |
| $\tau_{00}$ ID | 0.03 | | |
| $\tau_{11}$ ID.Intensity2 | 0.02 | | |
| $\tau_{11}$ ID.Intensity3 | 0.02 | | |
| $\rho_{01}$ | 0.16 | | |
|  | -0.15 |  |  |
| ICC | 0.22 |  |  |
| N<sub>ID</sub> | 21 |  |  |
| Observations | 15033 |  |  |
| Marginal R<sup>2</sup> / Conditional R<sup>2</sup> | 0.231 / 0.396 |  |  |

## Discussion

This study shows that cutaneous stimulation of peripheral nerves in the lower limb can evoke reflex responses in the PFM in uninjured, healthy adults. Bilateral tibial nerve stimulation evoked larger reflex responses compared to unilateral stimulation. PFM reflex amplitudes increased with the intensity of tibial nerve stimulation but not with superficial peroneal nerve stimulation, indicating a nerve-specific modulation of PFM activity. In most conditions in most participants, PFM responses were observed in the absence of responses in the GM, suggesting the reflex response observed in PFM is less likely from cross-talk.

### Neural connection between the PFM and the lower limb

In the current study, the reflex responses observed in PFM suggest a neural connection between lower limb afferents and muscular structures in the pelvic region. Analogous neural connections have been observed in animal models using urological or neurological measurements (Buss and Shefchyk 1999; Gad et al. 2016; McPherson 1966; Morrison et al. 1995; Sato et al. 1980; Yecies et al. 2018). In anesthetized cats, large rhythmic fluctuations in micturition bladder pressure (a sign of overactive bladder) were inhibited after stimulation of the superficial and deep peroneal nerve (Sato et al. 1980). In precollicular-postmamillary decerebrate cats, primary afferent depolarization was seen in the urethral afferents following stimulation of the sural and femoral cutaneous nerve, suggesting a potential for inducing an excitatory post-synaptic potentiation of motoneurons of the urethral muscle without descending inputs (Buss and Shefchyk 1999). In unanesthetized, spinalized rats, stimulation to either the tibial nerve or pudendal nerve evoked similar recruitment curves (i.e. input-output relationships) in the external urethral muscles, which also suggests an integrated neural network controlling bladder function and lower limb function (Gad et al. 2016).

Few studies have systematically investigated such reflex connections in human participants. Responses in the sphincter muscle have been reported in adults both with or without spinal lesions following tibial nerve stimulation (Pedersen et al. 1978) as well as in children with spinal cord injury following pinprick to the foot sole (Hallett et al. 1984). There is also some clinical evidence supporting the effect of tibial nerve stimulation on structures controlling bladder function. For example, in traditional Chinese medicine, acupuncture points in the lower limb have been used to treat bladder symptoms (Bergström et al. 2000; Lin et al. 2020). Transcutaneous or percutaneous electrical nerve stimulation of the tibial nerve is also prescribed clinically to treat overactive bladder(Booth et al. 2018; Wall and Heesakkers 2017).

In the current study, we observed a facilitatory reflex response in the PFM with the onset latency at 53.5 ± 16.7 ms and peak latency at 113.0 ± 19.1 ms following tibial nerve stimulation. The latency of tibial nerve stimulation-evoked PFM response has only been reported in one study based on our search. In this study (Pedersen et al. 1978), reflex in the anal sphincter was measured through needle electrodes and was seen at 93 ± 21.1 ms after tibial nerve stimulation(Pedersen et al. 1978). The discrepancy may be due to different electrode placement or different approaches to determining the reflex onset, as it was not reported in the previous study. Our findings provide neurological evidence showing the potential mechanisms underlying the therapeutic effects of tibial nerve stimulation on bladder overactivity in humans.

### Enhanced sensory inputs from the bottom of the feet amplify PFM reflex

In unilateral and bilateral tibial nerve stimulation conditions, PFM reflex amplitude increased with stimulation intensity. Larger PFM reflex responses were evoked by bilateral tibial nerve stimulation compared to unilateral stimulation, suggesting enhanced sensory inputs from the bottom of the feet can modulate the neural connections between the lower limb and pelvic structure.

Sensory feedback from the foot soles conveys information about the terrain, which is critical in modulating leg muscle activity and guiding foot placement during walking and other locomotor tasks (Hiebert and Pearson 1999; Zehr et al. 1997). This modulatory effect was indirectly observed in PFM during walking. Williams et al. (Williams et al. 2022) measured PFM activity during walking and jogging in healthy adults and observed greater activity that tended to align with the mid-stance to push-off phase. When comparing the PFM activity during supported walking in people with spinal cord injury, greater PFM activity was seen when walking in an exoskeleton that requires self-initiated weight shifting (Ekso) compared to completely passive walking with full-body weight support (Lokomat) (Zhou 2023). Both studies showed increased PFM activity when there is greater weight-bearing in the lower limb. However, because greater IAP also corresponds to the stance phase of walking (Grillner et al. 1978), the contribution of lower limb loading vs. IAP on PFM activity could not be distinguished in these previous studies.

In the current study, greater PFM response at higher stimulation intensities and with bilateral tibial nerve stimulation indicates that sensory inputs from the bottom of the feet can amplify the excitability of the interneurons that mediate the neural connection between PFM and lower limb muscles. Such an amplification effect was also observed during arm-leg cycling when the upper and lower limbs were stimulated simultaneously vs. individually (Nakajima et al. 2013). Nakajima et al. (Nakajima et al. 2013) found that simultaneous stimulation of the superficial radial nerve at the wrist and the SP nerve at the ankle evoked significantly larger reflexes in the vastus lateralis compared to the mathematical summation of the reflexes evoked by stimulating each nerve alone. Because sensory inputs from the skin surface of the hand and feet convey information about the obstacles and impediments during locomotive tasks, the authors suggested that convergent sensory inputs from the task-relevant limbs amplified the somatosensory linkage between limbs, which are important in priming for stumble correction and falling preparation (Nakajima et al. 2013). In our study, the influence of IAP changes on PFM activity is minimized since all conditions were completed during quiet standing. Increased reflex amplitude from low to high intensity and from Tib-R to Tib-B conditions highlights the modulation effects of the sensory inputs from the bottom of the feet.

### Nerve-specific responses and functional relevance

We found that the PFM reflex amplitude modulated with tibial nerve stimulation intensity, but a similar modulation was not observed with superficial peroneal nerve stimulation. Such nerve-specific cutaneous reflex modulation is also well known from the literature of reflex modulation during locomotion. The effects of cutaneous inputs on leg muscle activity during walking have been well studied in both animals and humans (Bouyer and Rossignol 2003a, 2003b; Forssberg 1979; Hiebert and Pearson 1999; Zehr et al. 1997, 1998b). Nerve-specific cutaneous reflex responses are seen in the leg muscles following sensory inputs from either the top or bottom of the foot. For instance, SP nerve stimulation during the swing phase of walking elicited an inhibitory response in the TA muscle, which was correlated with reduced ankle dorsiflexion. In contrast, tibial nerve stimulation evoked an inhibitory reflex in the TA muscles at the late swing phase but a facilitatory reflex at the stance-to-swing transition (Zehr et al. 1997). These nerve-specific modulations allow for flexible responses in muscles that are adaptable to the the phase of locomotion.

Nerve-specific responses are also seen in the neuro-urological studies in animal models. In neurologically intact, anesthetized cats, as well as cat models of bladder overactivity, 30 minutes of continuous tibial nerve stimulation resulted in an increase in bladder capacity by 44% (Park et al. 2020). Bladder infusion with acetic acid reduces the bladder capacity to around 20% of the saline-infusion control trial. However, tibial nerve stimulation during bladder infusion with acetic acid diminished the effects of acetic acid and increased bladder capacity to ∼40-60% of the control (Tai et al. 2011b, 2011a). In contrast, peroneal nerve stimulation has been used to treat bladder underactivity and nonobstructive urinary retention. Peroneal nerve stimulation applied during cystography increased the bladder contraction amplitude from 55.9 ± 5.0% to ∼70% of control (Chen et al. 2021), and increased voiding efficiency from 49.5 ± 16.8% to over 70% of control (Chen et al. 2021, 2022). These studies suggest sensory inputs from the top and bottom of the feet have opposite effects on the muscular structure in the pelvic region and bladder function.

We propose that the functional significance of the interaction between different sensory inputs from the lower limb and pelvic structures may be related to the influence of different postures on the synergy between PFM and detrusor muscle during bladder filling vs. voiding. As the bladder fills, detrusor muscle relaxation supports urine storage capacity and regulation of bladder pressure. Throughout, the PFM must remain activated to ensure continence in spite of potential IAP changes associated with posture or task transitions. Afferent feedback from the foot soles (e.g. via the distal tibial nerve) is well suited to monitor such critical changes in IAP associated with motor tasks, thereby giving rise to a facilitatory reflex pathway to the PFM that would support continence. In contrast, efficient voiding requires detrusor contraction accompanied by relaxation of the PFM. Considering that voiding usually occurs in a squatting position, sensory inputs associated with flexion may convey the information of “posture readiness” for voiding. This postural modulation was reported in the external urethral sphincters (EUS) of cats with spinal transection (Jolesz et al. 1982) where the normally facilitatory EUS reflexes evoked by subcutaneous stimulation to the lateral side of the hindlimb were altered depending on whether the limb was placed in flexion or extension. Unilateral reflexes recorded from the left nerve to the EUS were inhibited when the ipsilateral hindlimb was flexed and contralateral limb was extended, but not when the ipsilateral side was extended and the contralateral limb was flexed. This inhibitory effect increased with increasing contrast in bilateral limb positioning, with almost complete suppression of the EUS reflex with full flexion of the ipsilateral hindlimb and full extension of the contralateral side (Jolesz et al. 1982). In our study, although we did not alter limb posture, the contrasting reflex effects of superficial peroneal vs. tibial nerve stimulation on the PFM suggests that the PFM that is more responsive to sensory inputs associated with leg extension (i.e. plantar surface of the foot) compared to limb flexion (dorsal surface of the foot). Future experiments in humans may be designed to further explore a potential modulatory role of limb position interacting with the nerve-specific effects we observed here.

### Methodological considerations

In the current study, all the tests were completed during a natural standing position. To ensure participants are comfortable and avoid possible confounding effects, we asked participants to empty their bladders before data collection and avoid drinking water during data collection. Since the PFM reflexes were measured at the early phase of bladder filling, inputs from the top of the feet may be less relevant to the task requirement of our study, as we discussed in the previous section. Due to the limitations of our experimental setup, we were not able to obtain a quantitative measurement of the bladder filling status, whether SP nerve stimulation can modulate PFM activity in different postures and/or bladder status needs further study.

### Clinical Implication

Incontinence, neurogenic detrusor overactivity, and other lower urinary tract dysfunctions affect over 60% of adults over 40 years old (Chapple et al. 2017; Coyne et al. 2009), and over 80% of people with spinal cord injury or other types of neurological conditions (Agrawal and Joshi 2015; Taweel and Seyam 2015). PFM exercises are usually the first line of non-invasive treatment to manage urinary incontinence (Bø 2004, 2012; McClurg et al. 2006), but because they depend on voluntary contractions, such interventions will be challenging for clinical populations with neurogenic lower urinary tract dysfunction. Our study shows that PFM activity can be modulated by sensory inputs from the lower limb. It provides a neurological foundation to explore accessible approaches, such as weight-bearing exercise or transcutaneous stimulation of the lower limb, to help facilitate the activation of the PFM and support the management of neurogenic bladder symptoms in populations who cannot voluntarily activate this muscle group.

## DATA AVAILABILITY

The source data are available to verified researchers upon request by contacting the corresponding author.

### ACKNOWLEDGEMENTS

We thank Amy Schneeberg for her support in the statistical analysis and Ms. Alison William for her technical assistance during data collection.

## DISCLOUSRE

No conflicts of interest, financial or otherwise, are declared by the authors.

## AUTHOR CONTRIBUTIONS

YS & TL conceived and designed research; YS & CH performed experiments; YS & CH analyzed data; YS interpreted results of experiments, prepared figures, drafted manuscript; YS, JFY, EPZ and TL edited and revised manuscript; YS and TL approved final version of the manuscript

